# What’s in Your Body of Water? Reducing the Psychological Distance of Pharmaceutical Pollution through Metaphoric Framing in Risk Communication (A Pilot Study)

**DOI:** 10.1101/575639

**Authors:** Alexandra Z. Millarhouse, Christine Vatovec, Meredith T. Niles, Adrian Ivakhiv

## Abstract

Aquatic pharmaceutical pollution poses ecotoxicological risks to the environment and human health. Consumer behaviors represent a significant source of pharmaceutical compounds found in water. Thus, understanding public perceptions of aquatic pharmaceutical pollution and developing effective risk communication techniques are critical to engaging society in the type of widespread change necessary for addressing the presence of pharmaceuticals in water. This mixed-methods pilot study applies conceptual metaphor theory in conjunction with construal level theory of psychological distance to assess the relationship of metaphoric framing to perceptions of aquatic pharmaceutical contamination across four principal dimensions of psychological distance (geographic, social, and temporal distance, and uncertainty). Additionally, this study assesses the direct impact of metaphor use on concern and willingness to act, which are positively associated with perceived psychological distance. Data were collected from a convenience sample (n = 20) of university students in Burlington, Vermont using cognitive interviewing. Results indicate that participants initially perceived pharmaceutical pollution as socially and geographically distant, temporally both distant and proximal, and agreed that the issue is certain. Representing aquatic pharmaceutical contamination through metaphor significantly reduced perceived social and geographic distance, suggesting a relationship between metaphoric framing and psychological distance warranting additional research. Metaphor use did not directly nor significantly impact concern or willingness to act. Additionally, participants preferred the metaphorically-framed visual intervention to the non-metaphor visual intervention. Theoretical and practical implications of metaphor use in risk communications are discussed.

## 1. INTRODUCTION

Pharmaceuticals are considered chemicals of emerging concern because of their ecotoxicological impacts on the environment and human health ^(1)^. As commercial chemicals, pharmaceuticals flow from consumers to the environment during their life cycle on a continual basis ^(2)^. Consumer behaviors, such as disposal of household medications (e.g. via the trash or down the drain), significantly contribute to the volume of pharmaceutical compounds found in water. For example, Dohle et al. ^(3)^ found that many people believe flushing drugs down the drain or toilet is unlikely to have harmful environmental impacts, particularly when the drugs are familiar over-the-counter (OTC) medications like pain relievers. And yet, as the authors point out, common pain relievers are one of the most frequently detected classes of pharmaceutical chemicals in the aquatic environment and can have severe adverse ecological impacts. Thus, understanding public perceptions of aquatic pharmaceutical pollution and developing effective risk communication techniques are critical to engaging society in the type of widespread change necessary for addressing the presence of pharmaceuticals in water. In this study, we apply psychological distance to characterize perceptions, attitudes and behaviors towards aquatic pharmaceutical contamination and conceptual metaphor theory to assess the impact of metaphor use in risk communication on relevant perceptions, attitudes and behaviors.

### 1.1 Pharmaceuticals in the Environment

Nationally, a growing body of literature documents the presence of pharmaceutical compounds in ground water ^(4)^, surface waters ^(5–7)^, and drinking water ^(8–10)^. In addition, pharmaceutical compounds have been detected in multiple aquatic species ^(11, 12)^, including edible species ^(13)^; and have been shown to cause reproductive and behavioral impacts in fish ^(14)^, bivalves ^(13)^ and zooplankton ^(15)^.

Consumers are a primary source of pharmaceuticals in the environment. Excretion, disposal, and bathing off topical medications are the main consumer routes by which pharmaceuticals enter the environment ^(16)^. As pharmaceutical use continues to rise, so does the volume of medications that may eventually enter the waste stream. Common household drug disposal methods, such as via municipal trash or household drains, lead to drinking and surface water contamination through landfill leachate and wastewater effluent ^(16)^. To reduce this preventable source of aquatic pharmaceutical contamination, government agencies, hospitals, pharmacies, and not-for-profits are now offering drug collection (“take-back”) programs as an alternative disposal method.

Although Americans are increasingly aware of aquatic pharmaceutical pollution and its consequences to human and environmental health, people continue to improperly store or dispose of medications ^(17)^ and many collection programs are not attracting significant participation. A recent study of university students indicated that in the last 12 months, a majority had purchased and used OTC (87%) and prescription drugs (77%) and had leftover medications of which they had not yet disposed ^(18)^. Of those who disposed of leftover drugs within the last year, only 1% with leftover OTC and <1% with leftover prescription medications did so through an environmentally-preferred drug take-back program.

Promoting widespread participation in drug collection programs is a useful first step in addressing aquatic pharmaceutical pollution ^(2)^. These initiatives encourage individual action and consumer responsibility, critical foundations for systems-level change ^(19, 20)^ to significantly reduce the presence of pharmaceutical chemicals in water. This study characterizes public perceptions and theoretical relationships between psychological distance and metaphor use to inform effective risk communication techniques for drug collection programs.

## 2. THEORETICAL GROUNDING AND APPROACHES

As cognitive frameworks, psychological distance and conceptual metaphor theory share a foundation that people experience and represent stimuli either as concrete or abstract, which impacts attitudes and behaviors ^(21)^. Psychological distance, an index of how near or far a concept is from a perceiver’s immediate experience, suggests a psychologically distant concept is represented through its abstract qualities (e.g. decontextualized features) and a psychologically close concept is construed in concrete terms (e.g. specific, perceptual details). Relevant attitudes and behavior are positively associated with psychological distance, and different distances (near or far) lead to different attitudes and behaviors. Conceptual metaphor theory suggests that people use metaphor as a cognitive tool to understand abstract concepts through more concrete terms (e.g. “life is a journey”). Metaphor use impacts people’s practical judgments of a target concept based on understood features of the source concept.

This observation has inspired a small but growing body of research that explores the theoretical and practical interactions between the two frameworks. However, studies have so far only investigated whether manipulating conditions of psychological distance impacts conceptual metaphor use. For example, research has shown that people are more likely to rely on metaphor when concepts are framed as psychologically distant (and abstract) versus near (and concrete)^(22)^. No one has yet examined whether metaphor use impacts perceived psychological distance, or relevant cognitive judgments such as attitudes and behavioral intentions.

This study addresses these gaps in the theoretical literature while also addressing the need to better understand public perceptions of aquatic pharmaceutical contamination. The objectives of this pilot study are (1) to assess the impact of metaphoric framing on perceived distance of the environmental hazard across dimensions (temporal, geographic, social group and uncertainty) (2) to assess the impact of metaphoric framing on concern for the environmental hazard across dimensions and (3) to assess the impact of metaphoric framing on willingness to act (Figure 1). This research contributes to the theoretical advancement of psychological distance and metaphor theories and informs practical risk communication strategies encouraging participation in drug take-back initiatives.

**Figure 1.**
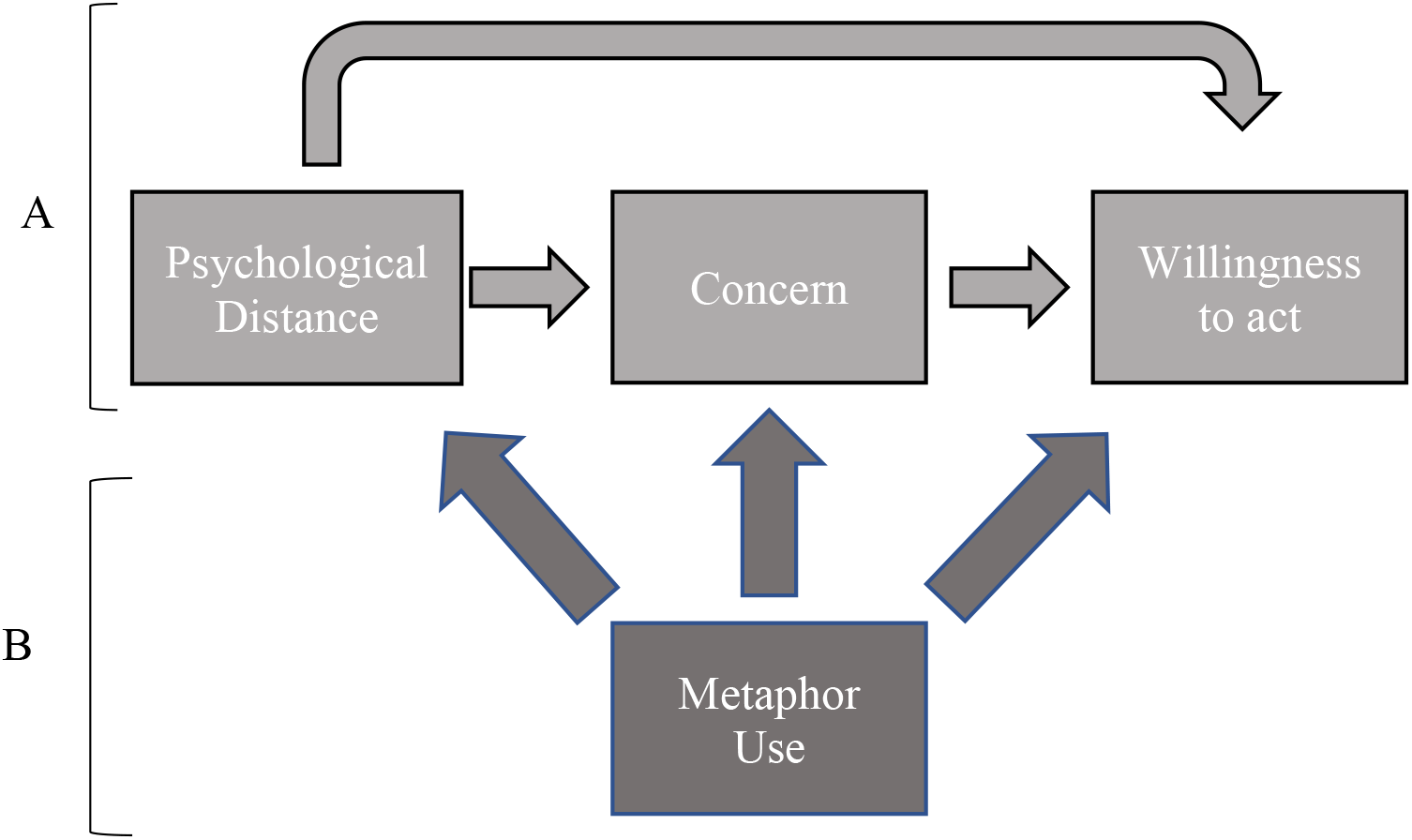
A simplified representation of the interplay between theoretical foundations and study objectives. A) An interpretation of the relationships between psychological distance and relevant attitudes and behavioral intentions based on findings from research applying psychological distance to environmental issues. Adopted from Spence ^(23)^; Niles ^(24)^. B) The relationships between metaphor use and psychological distance assessed in the objectives of the present study.

### 2.1 Construal level theory and psychological distance

Construal level theory ^(25, 26)^ posits that people perceive events, objects, actions, and other stimuli either as low-level (understood in specific terms) or high-level (conceptualized through global terms) constructs, which are inextricably linked to psychological distance. Within construal level theory, psychological distance is the mental distance perceived between a stimulus and the perceiver’s direct experience of their self in the present moment ^(27)^. Psychologically close stimuli tend to be low-level construals, understood through sensory and/or concrete terms ^(27)^. Psychologically distant stimuli are generally high-level construals understood through abstract, global terms ^(25)^.

Psychological distance is frequently studied through four primary dimensions: uncertainty, social group, geography and time. An event is psychologically closer when it is more likely to occur (uncertainty), happens to people like oneself (social group), occurs nearby (geographic) and takes place in the present or near future/past (time) ^(28)^. Psychologically distant events are perceived as unlikely to occur, happening to people unlike oneself, occurring far away and taking place in the distant future/past. Experimental evidence suggests that the dimensions are positively associated, so thinking about one dimension in psychologically close or distant terms may impact the cognitive processing of other dimensions (e.g., thinking about people unlike oneself may prime one to perceive a greater geographic distance) ^(27)^.

Psychological distance and construal level theory have wide-ranging implications for understanding and motivating human thought and behavior. Research has shown that when a concept is perceived as psychologically distant, people make choices based on their values (i.e. kindness); when something is represented as psychologically close, specific, contextual details like feasibility concerns (e.g. expected time commitment) and anticipated outcomes guide decisions ^(26)^. Additionally, the perceived distance of a target motivates people to different kinds of behaviors ^(29)^ and attitudes ^(28)^. For example, exploring the effects of psychological distance on farmers’ intentions to adopt different types of behaviors in response to climate change, Haden et al. ^(29)^ found that psychologically distant concerns impact farmers’ likelihood of adopting climate change *mitigation* practices (i.e. buying fuel efficient farm equipment) with abstract implications; while the intention to adopt climate change *adaptation* practices is influenced by feasibility concerns connected to psychological closeness (e.g. local water availability).

Consistent with other experimental studies of psychological distance, these studies demonstrate that related attitudes and behaviors are impacted by perceptions of psychological distance and suggest risk communication should intentionally and effectively frame psychological distance to produce desired responses to specific environmental issues ^(23)^. Specifically, framing risk communications to reduce the perceived psychological distance of a target issue may promote concern and intent to act ^(30)^.

### 2.2 Conceptual metaphor theory

Conceptual metaphor theory states that people rely on metaphors as a cognitive tool to make sense of abstract concepts through more concrete terms ^(31, 32)^. Metaphors in this context are conventional, everyday metaphors used by regular people ^(33)^. According to Geary ^(34)^, English speakers typically use about one metaphor for every 10-25 words spoken, or about six metaphors per minute ^(21)^.

In the metaphor framing model, metaphoric description (“using terms from another domain to talk about an event”) primes metaphoric encoding (“using schemas from another domain to think about an event”) ^(33)^. This results in the perceiver transferring knowledge of a source concept to interpret a target concept ^(22)^. Typically, source concepts are more easily comprehended and concrete experiences, whereas target concepts tend to be complex, hard to understand and more abstract ^(21)^. For example, past research demonstrates that metaphorically evoking the experience of protecting one’s body from contamination impacts people’s judgments about their country’s immigration policy. In two different studies, Americans more frequently opposed open immigration policies after being motivated to protect their own bodies from harmful (versus neutral) fictional bacteria ^(35, 22)^.

Exposure to different metaphors produces different effects on a person’s practical judgments. For example, investigating the consequences of stock market commentators’ use of metaphors on the judgments of investors, Morris et al. ^(33)^ found that exposing participants to agent-metaphors that implied an “enduring internal disposition” reflected through observed price trends (e.g. “The Nasdaq climbed higher”) resulted in an increased expectation that a present price trend would persist the next day. In contrast, neither object-metaphors, which do not imply an internal motivation (i.e. “The Nasdaq was pushed higher”), nor non-metaphorical descriptions of the stock market impacted investors’ expectations of trend continuance.

Importantly, certain conditions are necessary in order for a metaphor to be activated and made useful as a conceptual tool. For example, a metaphor needs to be culturally and contextually relevant ^(36)^. It also needs to be accessible to the individual perceiver and aligned with their unique epistemological and ontological perspectives. Steen et al. ^(37)^ suggest that reinforcing the metaphor through additional supportive textual/contextual references increases metaphoric transfer. Recent research also indicates that certain conditions of psychological distance may also be required for metaphoric activation ^(22)^.

## 3. METHODS

In this pilot study, we applied a mixed-methods approach to characterize perceptions of psychological distance, concern and behavioral intentions towards aquatic pharmaceutical contamination and whether metaphor use in risk communications impacts these perceptions. The study was approved by the University of Vermont (UVM) Institutional Review Board.

Data collection took place in Burlington, Vermont, between September 20 and November 7, 2016. All currently enrolled students (over the age of 18) able to meet in person on the UVM campus were eligible. The tailored design method ^(38)^ was applied to all phases of the study. Volunteer participants were recruited through email announcements sent through student listservs. Confidential, individual in-person interviews were audio-recorded and lasted 55 minutes on average.

### 3.1 Sampling procedure

To understand whether metaphor use impacts perceptions of psychological distance, concern and behavioral intentions, the study was designed as a crossover study to reduce potential order and performance variation effects (e.g. practice, boredom, fatigue, etc.). Participants were randomly assigned a treatment sequence group (group A or group B), counterbalancing the order of metaphor and non-metaphor treatments. Each treatment group was composed of half of the total sample (n = 10; see Table I).

**Table I.** Procedure.

| Interview Survey<br>Treatment<br>Order | Treatment<br>Group |  |
| --- | --- | --- |
|  | A (n = 10) | B (n = 10) |
| 1. Consent & 2.<br>Instructions | ✓ | ✓ |
| 3. Questions on<br>awareness of topic | ✓ | ✓ |
| 4. Psychological<br>distance survey | ✓ | ✓ |
| 5. 1<sup>st</sup> Visual | Metaphor<br>treatment | Non-metaphor<br>treatment |
| 6. Psychological<br>distance survey | ✓ | ✓ |
| 7. 2<sup>nd</sup> Visual | Non-metaphor<br>treatment | Metaphor<br>treatment |
| 8. Psychological<br>distance survey | ✓ | ✓ |
| 9. Questions on<br>current behavior | ✓ | ✓ |
| 10. New Ecological<br>Paradigm Scale<sup>(43)</sup> | ✓ | ✓ |
| 10. Demographic<br>information | ✓ | ✓ |

Data was collected using cognitive interviewing ^(39, 40)^, a semi-structured, interactive, and in-depth qualitative survey method ^(41)^, in which participants respond to a survey questionnaire while discussing aloud their thought processes and answer selections. Cognitive interviewing seeks to understand how respondents understand questions and the cognitive processes that are used to produce an answer ^(39)^ and requires a small but deliberate sample (typically 15 – 40 participants).

Participants were instructed to read each survey question aloud, select an answer, and discuss their thought processes with the interviewer through one of the six general but directed types of cognitive interviewing prompts outlined in Groves et al. ^(42)^.

Between- and within-group results were compared using descriptive statistical analysis and qualitative analysis. A Wilcoxon signed ranks test (assuming a null hypothesis of no change in mean response between pairs) was used to assess whether any within-group changes after the first and second visual treatments were statistically significant at p < 0.10. The decision to use a 10% significance was based on the small sample size and pilot-nature of this mixed-methods study.

It should be noted that following first treatment and subsequent repetition of the survey, most participants ascertained that we wanted to know if the visual changed how they thought about the issue. Consequently, the crossover study design did not successfully prevent order effects and people became practiced in the survey after the first treatment. As a result, the second treatment had an insignificant and unclear impact on both groups and only baseline and first treatment results are reported here.

### 3.2 Materials

#### 3.2.1 Survey instrument

The survey questionnaire was composed of (1) baseline perceptions of and behavioral intentions towards aquatic pharmaceutical contamination, (2) perceptions and behavioral intentions after viewing the first of two poster advertisements for safe drug disposal, (3) perceptions and behavioral intentions after viewing second poster advertisement for safe drug disposal, and (4) demographics (see Supplemental Materials for the full survey instrument). Survey questions assessed perceptions of distance and levels of concern across the four primary dimensions of distance (geographic, social group, uncertainty, and temporal) and were adapted from Spence et al. ^(23)^.

#### 3.2.2 Visual Treatments

Two fictional posters were developed as potential advertisements for drug collection programs, one framing the issue through a “nature as body” metaphor and one without this metaphor. The posters were identical in design and visual organization but differed in content (see Supplemental Materials for both visual treatments). The metaphor poster employed the root metaphor of “nature as body” to prime participants to protect their own bodies from contamination. Jackson ^(44)^ demonstrates that personal and nature “bodies” are metaphorically linked in many cultural and religious traditions.

## 5. RESULTS & DISCUSSION

Based on the Fall 2016 UVM Enrollment Report, the sample was roughly representative of the overall UVM student population in key demographic characteristics including gender, race, student level (undergraduate versus graduate), and in-state versus out-of-state residence ^(45)^. Survey respondents were 45% male and 50% female (5% of respondents did not select a sex). A majority of participants (85%) presently resided in Burlington, Vermont, identified their race as White/Caucasian (80%) and ethnicity as not Hispanic or Latino/a (100%), were undergraduate-level students (90%), and were out-of-state residents (65%). The sample was not representative of the UVM student population in undergraduate degree year or UVM school/college affiliation (see Supplemental Materials for full demographics).

This pilot study resulted in three primary findings. First, participants initially reported perceiving the issue of pharmaceutical pollution as distant across social and geographic dimensions, while perceiving the issue as comparable at near and far distances when considering the dimensions of time and uncertainty. Second, representing pharmaceutical pollution through the nature-as-body metaphor significantly reduced perceived social and geographic distance, as compared to the non-metaphor-based representation, but did not significantly impact perceived distance across temporal or uncertainty dimensions. Finally, the metaphor-based treatment did not significantly impact concern or behavioral intentions.

### 5.1 Initial perceptions of pharmaceutical pollution: psychological distance, concern, and willingness to act across dimensions

Participants of treatment groups A and B more strongly agreed that pharmaceutical pollution is a distant geographic and social issue (versus near), and expressed higher levels of concern for the issue at greater geographic and social distances (Table II). However, in response to questions regarding temporal distance and uncertainty, participants expressed nearly equal agreement that pharmaceutical pollution is a near and far issue. Unlike past research which suggests the four dimensions of distance are positively associated ^(27)^, our results indicate people may perceive varying levels of distance depending on the dimension.

The perception that the issue is more likely to impact other places and people may be due to spatial bias (environmental problems are believed to be worse at global versus local levels ^(46)^, especially by younger and happier people ^(47)^) and/or spatial optimism (environmental conditions are better here than elsewhere) ^(28, 48)^. For example, in a study assessing California farmers’ perceptions of climate change policy risks, Niles et al. ^(49)^ found that overall farmers believe climate change poses greater risks to agriculture globally (far) than to agriculture in Yolo County, California (near).

These cognitive biases have implications for behavior. Believing environmental problems to be more severe at a global level can lead to decreased feelings of self-efficacy (feeling able to do something about the problem) and responsibility for the problem ^(46)^, which in turn discourages public engagement. Likewise, our baseline results indicate people felt more motivated (a value-driven, high-level construal) than prepared (a low-level construal motivated by feasibility concerns) to participate in pharmaceutical take-back initiatives, which may be connected to perceptions that aquatic pharmaceutical contamination is a distant issue.

**Table II.** Initial perceptions of psychological distance, concern and willingness to act across dimensions for treatment groups A and B (n = 20).

| CONSTRUCT | DIMENSION<br>DISTANCE | QUESTION | RESPONSE<br>OPTIONS | INITIAL<br>MEAN<br>RESPONSE | %<br>UNSURE |
| --- | --- | --- | --- | --- | --- |
| DISTANCE | Geographic<br>Near | My local area is likely to be affected by the presence of pharmaceuticals in the water. |  | 3.3 | 0 |
|  | Geographic<br>Far | The presence of pharmaceuticals in the water will *mostly* affect areas that are far away from here. |  | 3.1 | 5 |
|  | Social<br>Near | People like me are likely to be affected by the presence of pharmaceuticals in the water. | 4-point scale<br>(4) Strongly Agree – (1) Strongly Disagree | 3.2 | 10 |
|  | Social<br>Far | Other people who are not like me are likely to be affected by the presence of pharmaceuticals in the water. |  | 3.5 | 5 |
|  | Uncertainty<br>Presence | Scientists are uncertain about the presence of pharmaceuticals in the water. |  | 1.7 | 10 |
|  | Uncertainty<br>Causes | Scientists are uncertain about what causes the presence of pharmaceuticals in the water. |  | 1.8 | 0 |
|  | Uncertainty<br>Problem | I am uncertain that the presence of pharmaceuticals in the water is really an issue. |  | 1.8 | 0 |
|  | Time<br>Near People | Do you think local residents will feel the effects of pharmaceuticals in the water? |  | 3.3 | 30 |
|  | Time<br>Far People | Do you think people in other areas around the world will feel the effects from the presence of pharmaceuticals in the water? | 4-point scale<br>(4) Yes, already feeling the effects – (1) No, I don't think the effects will be felt | 3.3 | 20 |
|  | Time<br>Near Environment | Do you think the local aquatic environment will feel the effects from the presence of pharmaceuticals in the water? |  | 4.0 | 10 |
|  | Time<br>Far Environment | Do you think aquatic environments in other places around the world will feel the effects from the presence of pharmaceuticals in the water? |  | 3.8 | 5 |
| **CONCERN** | Geographic<br>Near | When I think about my local area, I am concerned about the presence pharmaceuticals in the water. |  | 3.4 | 15 |
|  | Geographic<br>Far | When I think about areas around the world, I am concerned about the presence pharmaceuticals in the water. |  | 3.7 | 0 |
|  | Social<br>Near | When I think about people like me, I am concerned about the presence of pharmaceuticals in the water. |  | 3.2 | 5 |
|  | Social<br>Far | When I think about other people who are different from me, I am concerned about the presence pharmaceuticals in the water. | 4-point scale<br>(4) Strongly Agree – (1) Strongly Disagree | 3.5 | 5 |
|  | Uncertainty<br>Environment | It is uncertain if the presence of pharmaceuticals in the water will have any effects on the environment. |  | 1.4 | 5 |
|  | Uncertainty<br>People | It is uncertain if the presence of pharmaceuticals in the water will have any effects on people. |  | 1.8 | 5 |
|  | Time<br>Near | When I think about the near future, I am concerned about the presence pharmaceuticals in the water. |  | 3.4 | 10 |
|  | Time<br>Far | When I think about the distant future, I am concerned about the presence of pharmaceuticals in the water. |  | 3.4 | 10 |
| **BEHAVIORAL INTENTIONS** | Prepared | I feel prepared to participate in a pharmaceutical take back initiative. | 4-point scale<br>(4) Strongly Agree – (1) Strongly Disagree | 3.4 | 5 |
|  | Motivated | I feel motivated to participate in a pharmaceutical take back initiative. |  | 3.6 | 10 |

### 5.2 The impacts of metaphor-based framing on perceptions of pharmaceutical pollution

#### 5.2.1 Qualitative assessment of metaphor effectiveness and reception

Qualitative data capturing people’s responses to the metaphor-framed visual indicate the metaphor successfully provoked people to think about bodily exposure while interpreting the issue of aquatic pharmaceutical contamination. Further, the majority of respondents preferred the metaphor-based visual to communicate about drug take-back programs.

All participants were asked to describe their experience of each visual treatment, allowing us to assess the impact of the metaphor. Comparing the two potential advertisements, a majority of respondents (55%) stated a preference for the metaphor visual, 15% preferred the non-metaphor visual and 30% could not be determined. While viewing the metaphor treatment, most people (55%) described thinking about exposure to their bodies and linking that to thinking about pharmaceuticals in the water.

> *“Asking the question, ‘what’s in your body of water?’ makes you really wonder what’s in your body of water, like what’s going into your body? And then obviously having these pills in front of the lake makes you wonder again. […] so, you’re like ‘oh drugs in my body! That’s not a good thing*!’” (Participant T).
>
> “*What’s in your body of water? […] if you ask this I would probably think what is the mechanism of the medication – what is this medication going to cause in your body – what’s in your body of water*…” (Participant A).

Comparatively, while viewing the non-metaphor treatment, nearly everyone described their reaction to seeing the types and/or quantity of drugs represented. Various reactions included shock, disinterest and familiarity, among others. People also often commented on the headline question, “Got Drugs?”, which is used in advertisements for the U.S. Drug Enforcement Agency’s biannual National Take-Back Day. Many remarked that in a college environment, this may not be as attention-grabbing as it could be in other community settings.

#### 5.2.2 Impact of metaphor use on psychological distance

Metaphor use significantly reduced the perceived psychological distance of pharmaceutical pollution, compared to the non-metaphor intervention and baseline results. After viewing the metaphor-based visual, treatment group A participants perceived aquatic pharmaceutical contamination as significantly geographically (p = 0.083) and socially (p = 0.034) closer than was indicated by their baseline responses. Comparatively, treatment group B, who viewed the non-metaphor treatment, had no significant changes in perceived psychological distance (Figure 2).

**Figure 2.**
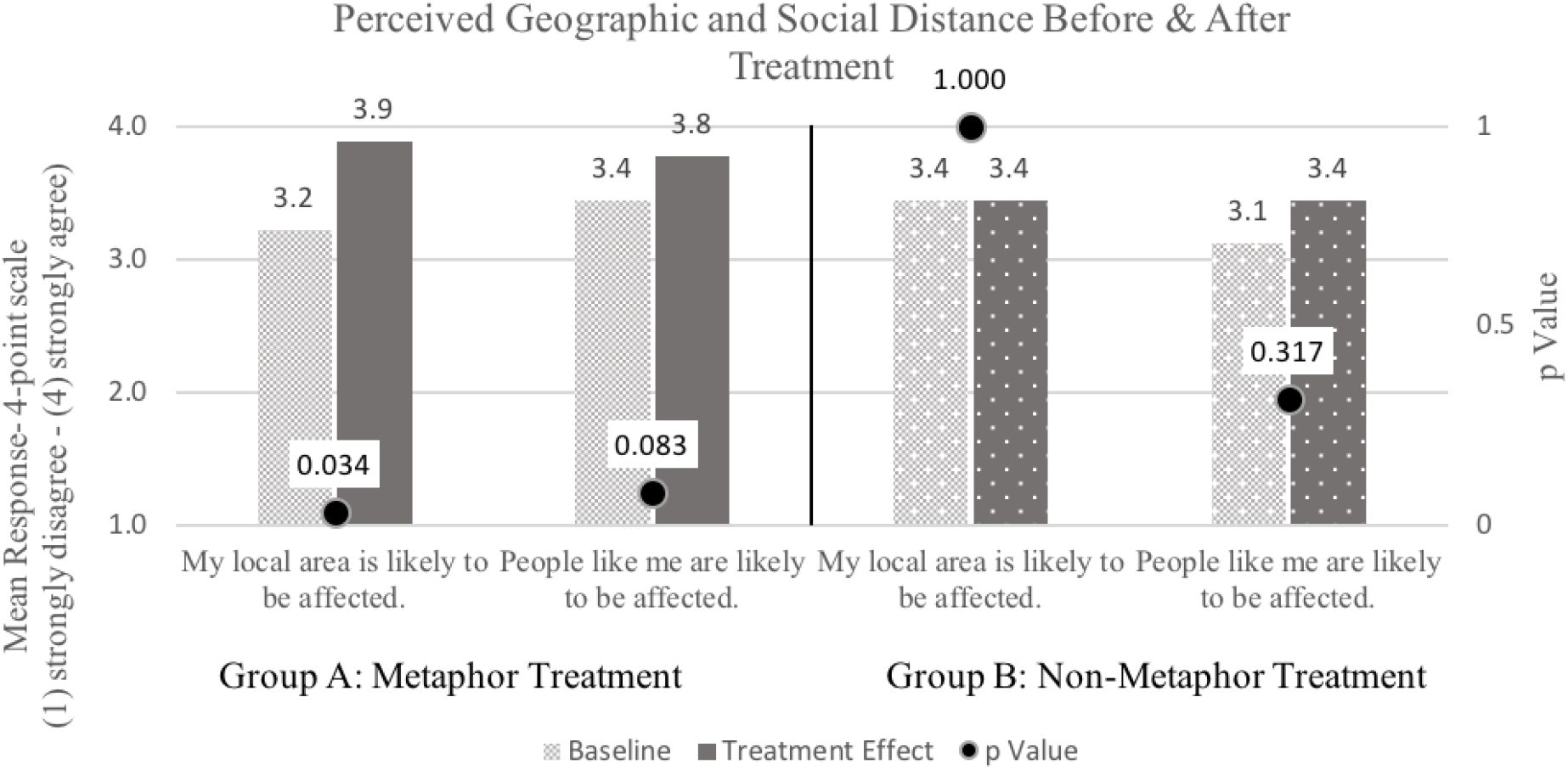
Change in perceived geographic and social distance for group A (n = 10) and group B (n = 10) between baseline and post-treatment.

Given the function of metaphor use (describing abstract concepts through concrete terms), we would expect metaphor-based framing to have a greater impact when conditions of psychological distance are present. In this study, initial responses indicated pharmaceutical pollution was only perceived as psychologically distant across geographic and social dimensions. Consistent with this expectation, metaphor use did not significantly alter perceived temporal distance or uncertainty as compared to baseline perceptions (Table III).

#### 5.2.3 Impact of metaphor use on concern and willingness to act

Representing the issue through metaphor had no direct, statistically significant effect on treatment group A’s initial levels of concern across dimensions, although overall concern increased across dimensions and distances. The non-metaphor treatment significantly increased treatment group B’s concern for geographically distant impacts (p = 0.083), compared with their baseline responses. In general, this treatment also increased concern across dimensions and distances, although no other change was statistically significant.

Metaphor use also had no direct, statistically significant impact on group A’s behavioral intentions (Figure 3), although people felt equally prepared and motivated to participate in a drug collection program (versus initially being more motivated than prepared). The non-metaphor visual also had no significant effect on group B’s behavioral intentions. People continued to feel more motivated than prepared.

**Figure 3.**
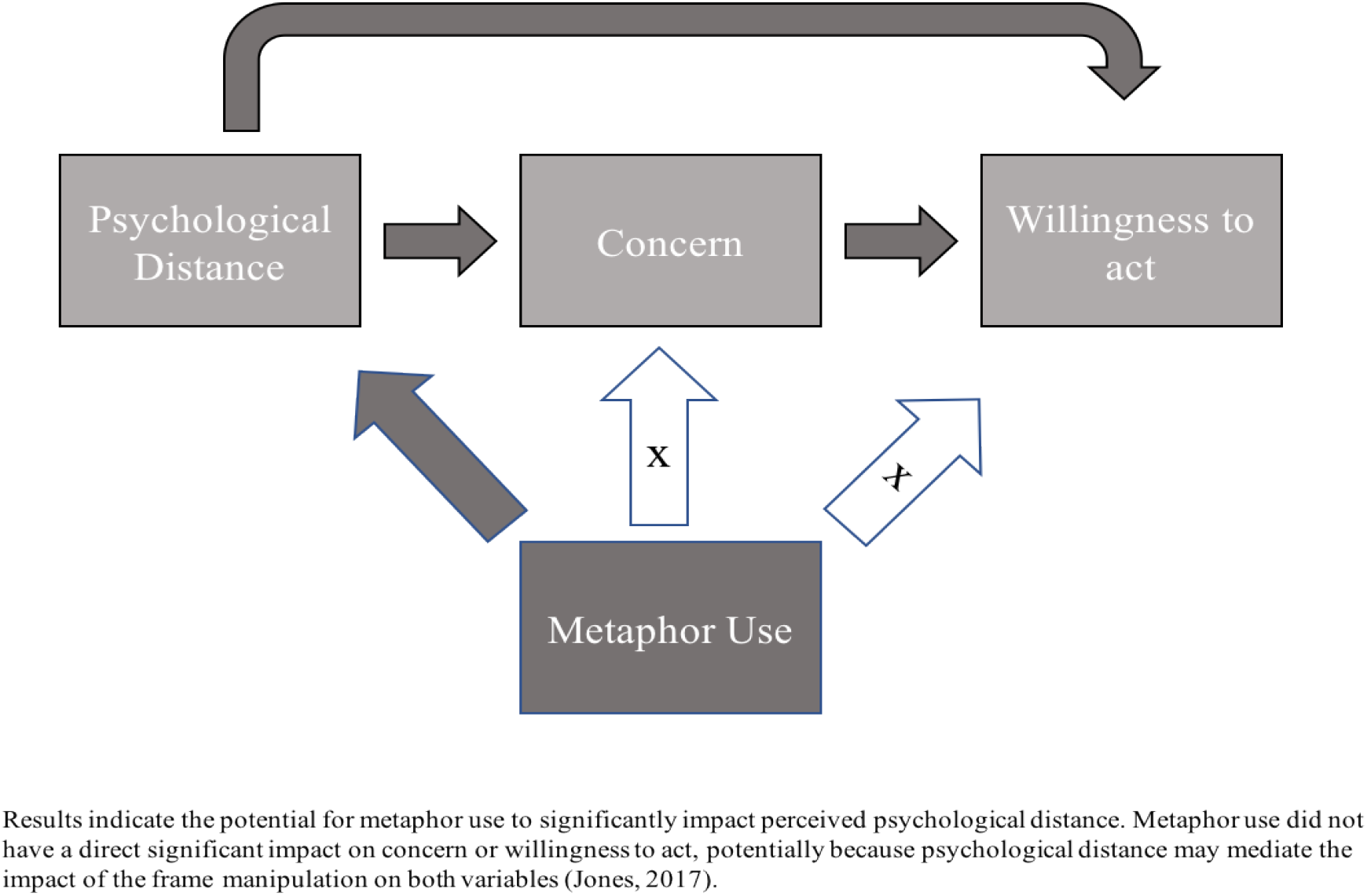
Observed and potential impacts of metaphor use on psychological distance, concern and willingness to act.

Recent research suggests psychological distance mediates the impact of message frame manipulations, like metaphoric framing, on concern and behavioral intentions. Jones et al. ^(30)^ found that framing messages to manipulate (reduce) psychological distance indirectly increased concern and willingness to act, but had no direct, statistically significant effect on either construct. Due to our small sample size, we could not assess whether psychological distance mediated the impact of metaphor use on concern and willingness to act; however, we strongly recommend that future research consider this particular relationship.

According to Rabinovich et al. ^(50)^, reducing psychological distance may be especially critical when specific individual actions are needed to achieve a relatively abstract goal, like participating in a drug collection program to reduce aquatic pharmaceutical contamination, which cannot be detected through the senses. Therefore, risk communication efforts to bring this issue closer may indirectly lead to greater concern and preparedness to act at an individual level.

**Table III.** Statistical results using Wilcoxon signed ranks test, a nonparametric method for analyzing differences and magnitude of difference between paired data that assumes a null hypothesis of zero difference ^(51, 52)^. Significant results (p < 0.10) are bolded for emphasis.

| CONSTRUCT | VARIABLE | METAPHOR<br>TREATMENT<br>(N = 10) |  | NON-METAPHOR<br>TREATMENT<br>(N = 10) |  |
| --- | --- | --- | --- | --- | --- |
|  |  | Z – Score | p Value* | Z – Score | p Value* |
| DISTANCE | Geographic Near | -2.121<sup>a</sup> | **0.034** | .000<sup>a</sup> | 1.000 |
|  | Geographic Far | -1.633<sup>b</sup> | 0.102 | -.816<sup>b</sup> | 0.414 |
|  | Social Near | -1.732<sup>a</sup> | **0.083** | -1.000<sup>b</sup> | 0.317 |
|  | Social Far | -.577<sup>b</sup> | 0.564 | -.447<sup>b</sup> | 0.655 |
|  | Uncertainty Presence | -1.342<sup>b</sup> | 0.180 | -1.000<sup>c</sup> | 0.317 |
|  | Uncertainty Causes | -1.000<sup>b</sup> | 0.317 | -1.342<sup>c</sup> | 0.180 |
|  | Uncertainty Problem | -.816<sup>a</sup> | 0.414 | -1.633<sup>c</sup> | 0.102 |
|  | Time Near People | -1.000<sup>b</sup> | 0.317 | -1.414<sup>b</sup> | 0.157 |
|  | Time Far People | -1.342<sup>a</sup> | 0.180 | -.577<sup>b</sup> | 0.564 |
|  | Time Near Environment | .000<sup>c</sup> | 1.000 | .000<sup>a</sup> | 1.000 |
|  | Time Far Environment | -1.000<sup>a</sup> | 0.317 | .000<sup>a</sup> | 1.000 |
| CONCERN | Geographic Near | -1.000<sup>a</sup> | 0.317 | -.816<sup>c</sup> | 0.414 |
|  | Geographic Far | .000<sup>c</sup> | 1.000 | -1.732<sup>b</sup> | **0.083** |
|  | Social Near | -1.000 <sup>a</sup> | 0.317 | -1.000 <sup>b</sup> | 0.317 |
|  | Social Far | -1.342 <sup>a</sup> | 0.180 | .000 <sup>a</sup> | 1.000 |
|  | Uncertainty Environment | -1.342 <sup>a</sup> | 0.180 | -.577 <sup>c</sup> | 0.564 |
|  | Uncertainty People | -1.633 <sup>b</sup> | 0.102 | -1.134 <sup>c</sup> | 0.257 |
|  | Time Near | -.816 <sup>a</sup> | 0.414 | -.577 <sup>c</sup> | 0.564 |
|  | Time Far | -1.342 <sup>a</sup> | 0.180 | -.577 <sup>b</sup> | 0.564 |
| **WILLINGNESS TO ACT** | Prepared | -1.000 <sup>a</sup> | 0.317 | .000 <sup>a</sup> | 1.000 |
|  | Motivated | -1.000 <sup>a</sup> | 0.317 | -.577 <sup>b</sup> | 0.564 |
|  | * statistically significant p values in bold.<br><sup>a</sup> Based on negative ranks.<br><sup>b</sup> Based on positive ranks.<br><sup>c</sup> The sum of negative ranks equals the sum of positive ranks. |  |  |  |  |

## 6. METHODOLOGICAL CONTRIBUTIONS AND LIMITATIONS

In this study, cognitive interviewing was used to understand how people perceive the issue of aquatic pharmaceutical contamination using questions adapted from Spence et al. ^(23)^ to measure psychological distance, concern and willingness to act. In the process of interviewing, we found key constructs were interpreted differently from person to person. For example, some people interpreted the geographic near construct, “my local area”, as the immediate area around Burlington, Vermont, while others assumed it meant their hometown located in another county, state or country. People often interpreted “near future” and “far future” as the future in general. Additionally, people commonly considered social factors when responding to questions assessing geographic distance and concern (e.g. regulations, environmental values, income, etc.) and likewise geographic features when answering questions assessing social distance and concern (e.g. proximity to water, physical location, etc.). These multiple interpretations could lead to inconsistent responses. As psychological distance becomes an increasingly popular framework for measuring perceptions, attitudes and behaviors, there is a need for standardized, validated language framing each dimension of distance that can be applied to studies across disciplines and topic areas.

We want to note that among this particular sample population it is possible that perceived psychological distance and concern for aquatic pharmaceutical contamination were impacted by (1) the proximity and visibility of Lake Champlain within Burlington, Vermont, ^(53)^ (2) prior awareness ^(54)^, (3) socially desirable responding (the tendency of questionnaire respondents to self-report socially acceptable answers) ^(55)^, and (4) the use of visual (versus text) communications, which may suggest proximity between a communicator and audience ^(56)^. Additionally, we know from qualitative data that answering questions about the topic in the context of a research study reduced perceived uncertainty about the issue and impacted people’s concerns for the near and far future. For example, some people felt less concerned about the far future because they assume that current studies, such as the one they were participating in, will lead to future solutions.

## 7. CONCLUSION

Consumer attitudes and behaviors significantly contribute to the presence of pharmaceutical chemicals in water systems with consequences to human and environmental health. In this study, we found that aquatic pharmaceutical contamination was initially perceived as geographically and socially distant. Further, compared to baseline perceptions and the non-metaphor treatment, using a nature-as-body metaphor to frame the issue of aquatic pharmaceutical contamination significantly reduced the perceived psychological distance of the issue, specifically across geographic and social dimensions of distance. While metaphor-based framing did not significantly impact participants’ concern or behavioral intentions, past research suggests that reducing perceived distance through a framing manipulation, like metaphor, may indirectly positively influence concern and preparedness to act. Finally, we found people interpret distances (near/far) and dimensions (geographic, social, temporal, and uncertainty) in different ways, suggesting the need for validated questions to consistently measure psychological distance.

While other studies have explored how framing psychological distance affects metaphor use, this study is the first that we know of to assess how metaphor use impacts perceived psychological distance. Our findings contribute to a growing body of theoretical literature exploring the utility of psychological distance and conceptual metaphor theory in understanding how people process and form cognitive judgements on everyday stimuli. Additionally, results from this study have practical applications for designing risk communications that effectively promote public engagement and action on the issue of aquatic pharmaceutical contamination.

## ACKNOWLEDGEMENTS

The authors thank the participants, Alan Howard, Director of the Statistical Counseling Clinic at the University of Vermont, for his assistance with statistical analysis and Jon Portman, Creative Director and Founding Partner at Oxbow Creative, for his graphic design input. Funding support for this project was generously provided by the Gund Institute for Environment, the Vermont Water Resources and Lake Studies Center, a USGS Water Resources Research Institute and the Rubenstein Graduate Student Association. This study was also supported in part by the University of Vermont Rubenstein School of Environment and Natural Resources, Office of Health Promotion Research, the College of Medicine Division of Hematology/Oncology, Department of Biochemistry, Department of Family Medicine, and Department of Surgery.

## REFERENCES

1. Environmental Protection Agency. White Paper Aquatic Life Criteria for Contaminants of Emerging Concern: Part I, General Challenges and Recommendations. In: OOEC Workgroup, editor., Series White Paper Aquatic Life Criteria for Contaminants of Emerging Concern: Part I, General Challenges and Recommendations. 2008.

2. Glassmeyer ST, Hinchey EK, Boehme SE et al. Disposal practices for unwanted residential medications in the United States. Environ Int, 2009; 35 (3):566–72.

3. Dohle S, Campbell VEA, Arvai JL. Consumer-perceived risks and choices about pharmaceuticals in the environment: a cross-sectional study. Environmental Health: A Global Access Science Source, 2013; 12:45.

4. Banzhaf S, Krein A, Scheytt T. Investigative approaches to determine exchange processes in the hyporheic zone of a low permeability riverbank. Hydrogeology Journal, 2011; 19 (3):591–601.

5. Kolpin DW, Furlong ET, Meyer MT et al. Pharmaceuticals, hormones, and other organic wastewater contaminants in U.S. streams, 1999-2000: a national reconnaissance. Environmental Science & Technology, 2002; 36 (6):1202.

6. Lara-Martín PA, Renfro AA, Cochran JK et al. Geochronologies of Pharmaceuticals in a Sewage-Impacted Estuarine Urban Setting (Jamaica Bay, New York). Environmental Science & Technology, 2015; 49 (10):5948–55.

7. Lara-Martín PA, González-Mazo E, Petrovic M et al. Occurrence, distribution and partitioning of nonionic surfactants and pharmaceuticals in the urbanized Long Island Sound Estuary (NY). Marine pollution bulletin, 2014; 85 (2):710–9.

8. Padhye LP, Yao H, Kung’u FT et al. Year-long evaluation on the occurrence and fate of pharmaceuticals, personal care products, and endocrine disrupting chemicals in an urban drinking water treatment plant. Water research, 2014; 51:266–76.

9. Focazio MJ, Kolpin DW, Barnes KK et al. A national reconnaissance for pharmaceuticals and other organic wastewater contaminants in the United States-II) Untreated drinking water sources. Science of the Total Environment, 2008; 402 (2–3):201–16.

10. Stackelberg PE, Gibs J, Furlong ET et al. Efficiency of conventional drinking-water-treatment processes in removal of pharmaceuticals and other organic compounds. Science of the Total Environment, 2007; 377 (2):255–72.

11. Brandao FP, Pereira JL, Goncalves F et al. The impact of paracetamol on selected biomarkers of the mollusc species Corbicula fluminea. Environmental Toxicology, 2014; 29 (1):74–83.

12. Ramirez AJ, Brain RA, Usenko S et al. Occurrence of Pharmaceuticals and Personal Care Products in Fish: Results of a National Pilot Study in the United States. Environmental Toxicology and Chemistry, 2009; 28 (12):2587–97.

13. Antunes S, Freitas R, Figueira E et al. Biochemical effects of acetaminophen in aquatic species: edible clams Venerupis decussata and Venerupis philippinarum. Environmental Science and Pollution Research, 2013; 20 (9):6658–66.

14. Jobling S, Williams R, Johnson A et al. Predicted exposures to steroid estrogens in U.K. rivers correlate with widespread sexual disruption in wild fish populations. Environmental Health Perspectives, 2006; 114 Suppl 1:32.

15. Flaherty CM, Dodson SI. Effects of pharmaceuticals on Daphnia survival, growth, and reproduction. Chemosphere, 2005; 61 (2):200–7.

16. Daughton C. Pharmaceuticals in the environment: sources and their management. Comprehensive analytical chemistry, 2007; 50:1–58.

17. Bound JP, Kitsou K, Voulvoulis N. Household disposal of pharmaceuticals and perception of risk to the environment. Environmental Toxicology and Pharmacology, 2006; 21 (3):301–7.

18. Vatovec C, Phillips P, Van Wagoner E et al. Investigating dynamic sources of pharmaceuticals: Demographic and seasonal use are more important than down-the-drain disposal in wastewater effluent in a University City setting. Science of the Total Environment, 2016; (572):906–14.

19. Daughton CG. Cradle-to-cradle stewardship of drugs for minimizing their environmental disposition while promoting human health. I. Rationale for and avenues toward a green pharmacy. Environmental Health Perspectives, 2003; 111 (5):757.

20. Daughton CG. Cradle-to-cradle stewardship of drugs for minimizing their environmental disposition while promoting human health. II. Drug disposal, waste reduction, and future directions. Environmental Health Perspectives, 2003; 111 (5):775.

21. Landau MJ, Robinson MD, Meier BP, editors. The Power of Metaphor: Examining its Influence on Social Life 1st edn. Washington, DC: American Psychological Association; 2014.

22. Jia L, Smith ER. Distance makes the metaphor grow stronger: A psychological distance model of metaphor use. Journal of Experimental Social Psychology, 2013; 49 (3):492.

23. Spence A, Poortinga W, Pidgeon N. The psychological distance of climate change. Risk Analysis, 2012; 32 (6):957–72.

24. Niles, M. T., Lubell, M., & Brown, M. (2015). How limiting factors drive agricultural adaptation to climate change. Agriculture, Ecosystems and Environment, 200, 178.

25. Liberman N, Förster J. Distancing from experienced self: How global-versus-local perception affects estimation of psychological distance. Journal of Personality and Social Psychology, 2009; 97 (2):203–16.

26. Trope Y, Liberman N. Construal-level theory of psychological distance. Psychological Review, 2010; 117 (2):440–63.

27. Bar-Anan Y, Liberman N, Trope Y et al. Automatic Processing of Psychological Distance: Evidence From a Stroop Task. Journal of Experimental Psychology: General, 2007; 136 (4):610–22.

28. Milfont TL, Abrahamse W, McCarthy N. Spatial and temporal biases in assessments of environmental conditions in New Zealand. New Zealand Journal of Psychology, 2011; 40 (2):56–67.

29. Haden VR, Niles MT, Lubell M et al. Global and Local Concerns: What Attitudes and Beliefs Motivate Farmers to Mitigate and Adapt to Climate Change? PLoS ONE, 2012; 7 (12):e52882.

30. Jones C, Hine DW, Marks AD. The future is now: reducing psychological distance to increase public engagement with climate change. Risk Analysis, 2016.

31. Gibbs RW. Why many concepts are metaphorical. Cognition, 1996; 61 (3):309–19.

32. Lakoff G, Johnson M. Metaphors We Live By Chicago: University of Chicago Press; 1980.

33. Morris MW, Sheldon OJ, Ames DR et al. Metaphors and the market: Consequences and preconditions of agent and object metaphors in stock market commentary. Organizational behavior and human decision processes, 2007; 102 (2):174–92.

34. Geary J. I is an other: the secret life of metaphor and how it shapes the way we see the world. 1st ed. New York: HarperCollins; 2011.

35. Landau MJ, Sullivan D, Greenberg J. Evidence that self-relevant motives and metaphoric framing interact to influence political and social attitudes. Psychological science, 2009; 20 (11):1421.

36. Landau MJ, Meier BP, Keefer LA. A Metaphor-Enriched Social Cognition. Psychological Bulletin, 2010; 136 (6):1045–67.

37. Steen GJ, Reijnierse WG, Burgers C. When Do Natural Language Metaphors Influence Reasoning? A Follow-Up Study to Thibodeau and Boroditsky (2013). PLoS ONE, 2014; 9 (12):e113536.

38. Dillman DA. Internet, phone, mail, and mixed-mode surveys: the tailored design method 4th ed. JD Smyth; LM Christian; DA Dillman, editors. Hoboken, New Jersey: Wiley; 2014. (JD Smyth; LM Christian; DA Dillman editors. Surveys).

39. Beatty PC, Willis GB. Research Synthesis: The Practice of Cognitive Interviewing. Public Opinion Quarterly, 2007; 71 (2):287–311.

40. Willis GB. Cognitive interviewing: A tool for improving questionnare design. Thousand Oaks, CA: Sage Publications; 2004.

41. de Leeuw ED, Hox JJ, Dillman DA. International handbook of survey methodology. New York & London: Lawrence Erlbaum Associates; 2008.

42. Groves RM, Fowler Jr FJ, Couper MP et al. Survey methodology: John Wiley & Sons; 2011.

43. Dunlap RE, Van Liere KD, Mertig AG et al. New Trends in Measuring Environmental Attitudes: Measuring Endorsement of the New Ecological Paradigm: A Revised NEP Scale. Journal of Social Issues, 2000; 56 (3):425–42.

44. Jackson M. Thinking through the body: An essay on understanding metaphor. Social Analysis: The International Journal of Social and Cultural Practice, 1983; (14):127–49.

45. University of Vermont. Census Enrollment Report, Fall 2016. 2016.

46. Uzzell DL. The psycho-spatial dimension of global environmental problems. Journal of Environmental Psychology, 2000; 20 (4):307–18.

47. Schultz P, Milfont T, Chance R et al. Cross-Cultural Evidence for Spatial Bias in Beliefs About the Severity of Environmental Problems. Environment and Behavior, 2014; 46 (3):267.

48. Gifford R, Scannell L, Kormos C et al. Temporal pessimism and spatial optimism in environmental assessments: An 18-nation study. Journal of Environmental Psychology, 2009; 29 (1):1–12.

49. Niles MT, Lubell M, Haden VR. Perceptions and responses to climate policy risks among California farmers. Global Environmental Change, 2013; 23 (6):1752.

50. Rabinovich A, Morton TA, Postmes T et al. Think global, act local: The effect of goal and mindset specificity on willingness to donate to an environmental organization. Journal of Environmental Psychology, 2009; 29 (4):391–9.

51. McDonald JH. Handbook of Biological Statistics (3rd ed.). Baltimore, Maryland: Sparky House Publishing; 2014.

52. Whitley E, Ball J. Statistics review 6: Nonparametric methods. Critical Care, 2002; 6 (6):509–13.

53. Milfont TL, Evans L, Sibley CG et al. Proximity to Coast Is Linked to Climate Change Belief. PLoS ONE, 2014; 9 (7).

54. Milfont TL. The Interplay Between Knowledge, Perceived Efficacy, and Concern About Global Warming and Climate Change: A One-Year Longitudinal Study. Risk Analysis, 2012; 32 (6):1003–20.

55. Van de Mortel TF. Faking it: social desirability response bias in self-report research. The Australian Journal of Advanced Nursing, 2008; 25 (4):40.

56. Amit E, Wakslak C, Trope Y. The use of visual and verbal means of communication across psychological distance. Personality and Social Psychology Bulletin, 2013; 39 (1):43–56.

